# CRISPR associated enzymes are mislocalized to the cytoplasm in iPSC-derived neurons resulting in KRAB-specific degradation

**DOI:** 10.1101/2024.10.19.619045

**Authors:** Gregory Cajka, Matthew Liu, Ophir Shalem

## Abstract

The use of CRISPR-associated enzymes in iPSC-derived neurons for precise gene targeting and high-throughput gene perturbation screens offers great potential but presents unique challenges compared to dividing cell lines. CRISPRi screens in iPSC-derived neurons and glia have already been successful in relating gene function to neurological phenotypes; however, loss of dCas9-KRAB expression after differentiation has been observed by many labs and has been largely ascribed to transgene silencing after differentiation. Here, we investigated the expression levels of different CRISPR enzymes in iPSC and Ngn2-derived neurons using piggybac delivery. We found that the commonly used dCas9-KRAB (using the *KOX1* domain) displayed dramatic reduction in protein expression levels following neuronal differentiation, yet surprisingly, nCas9 constructs retained comparable protein expression between iPSCs and neurons. We further found that CRISPR constructs, primarily relying on the SV40 Nuclear Localization Signal (NLS), fail to efficiently localize to the nuclei of neurons, despite having robust nuclear levels in iPSCs, leading to KRAB-specific cytoplasmic degradation. By adding a neuronal-specific NLS, we were able to correct neuronal nuclear localization and protein expression, confirming the contribution of mislocalization to the instability of dCas9-KRAB in neurons. As the lack of nuclear localization can have a profound impact on editing and gene perturbation efficiency, we suggest further investigation across both cultured and in-vivo postmitotic cell models.

## Introduction

The ability to differentiate induced pluripotent stem cells (iPSCs) directly into human neurons introduced a powerful and easily accessible cell model to study the molecular biology of neurological diseases and evaluate potential therapeutic approaches(Pang et al. 2011). Efficient implementation of tools for genetic manipulation, such as CRISPR-associated enzymes for gene knockout, editing, and knockdown, is essential for this model to realize its full potential, yet such implementation is frequently associated with unique challenges compared to immortalized cell lines. For example, loss-of-function screens using double strand break induced frameshifts are widespread and have led to numerous discoveries in immortalized cell lines, yet their implementation in iPSCs has been challenging due to increased sensitivity to DNA damage (Ihry et al. 2018). CRISPR inhibition and activation has emerged as a powerful and orthogonal approach to CRISPR knockout, which could be utilized in iPSCs as it works by recruiting repression or activation proteins to gene promoters and does not require the induction of DNA damage for its activity(Tian et al. 2019, 2021).

CRISPR inhibition and activation screens in iPSC-derived neurons have been successful in uncovering genes associated with several neuronal phenotypes including, survival(Tian et al. 2019), sensitivity to stress(Tian et al. 2021), tau aggregation(Parra Bravo et al. 2024) and the regulation of alpha-synuclein protein levels(Santhosh Kumar et al. 2024). In all of these cases, the screens have relied on introduction of genetic perturbations at the iPSC or early neuronal precursor stage, followed by evaluation of phenotypes after differentiation into mature neurons. This can confound phenotypes related to neuronal maintenance with defects in differentiation and maturation. It may also limit our ability to study genes that are essential for iPSC survival and growth but with distinct roles in post-mitotic neurons. However, developing screening systems where genetic perturbations are performed directly in neurons faces several biological and technical hurdles that likewise severely limit the development of CRISPR-based therapeutics, which ideally require high efficiency at low doses.Differential stability of proteins and RNA and differential chromatin organization can make it challenging to predict and evaluate the effectiveness of perturbations. And the efficiency of these perturbations is dependent upon ensuring high nuclear expression levels of CRISPR machinery at the time of their introduction.

## Results

### dCas9-KRAB exhibits neuron-specific cytoplasmic mislocalization and KRAB-dependent protein depletion in iPSC-derived NGN2 neurons

To move toward developing efficient systems for inducible gene perturbations later into neuronal differentiation, we longitudinally tested the expression levels of two CRISPR cassettes expressed in stable iPSC lines generated using piggybac integration into KOLF2.1J iPSCs with NGN2 integration at the AAVS1 locus(Pantazis et al. 2022). The two donor vectors were identical apart from one expressing dCas9 with the *KOX1* KRAB domain and the other expressing nCas9 (Fig 1A). Both were based on previously published dCas9-KRAB constructs(Tian et al. 2019; Gilbert et al. 2014; Qi et al. 2013) used to engineer iPSCs with targeted integration. Polyclonal iPSC lines were generated using piggybac transfection and selection with blasticidin, followed by fluorescence cell sorting based on tagBFP expression levels. We then compared the expression levels of dCas9-KRAB and nCas9 with flow cytometry in iPSCs and in neurons 14 days post-induction of differentiation with NGN2 expression by treatment with doxycycline (Fig 1B). Surprisingly, we found that dCas9-KRAB and nCas9 showed dramatic differences in fluorescence post-differentiation. In iPSCs, both constructs demonstrated comparable fluorescence levels, but, in Day 14 neurons, dCas9-KRAB fluorescence was essentially undetectable while nCas9 fluorescence remained robust, with a distribution comparable to that in iPSCs.

**Figure 1.**
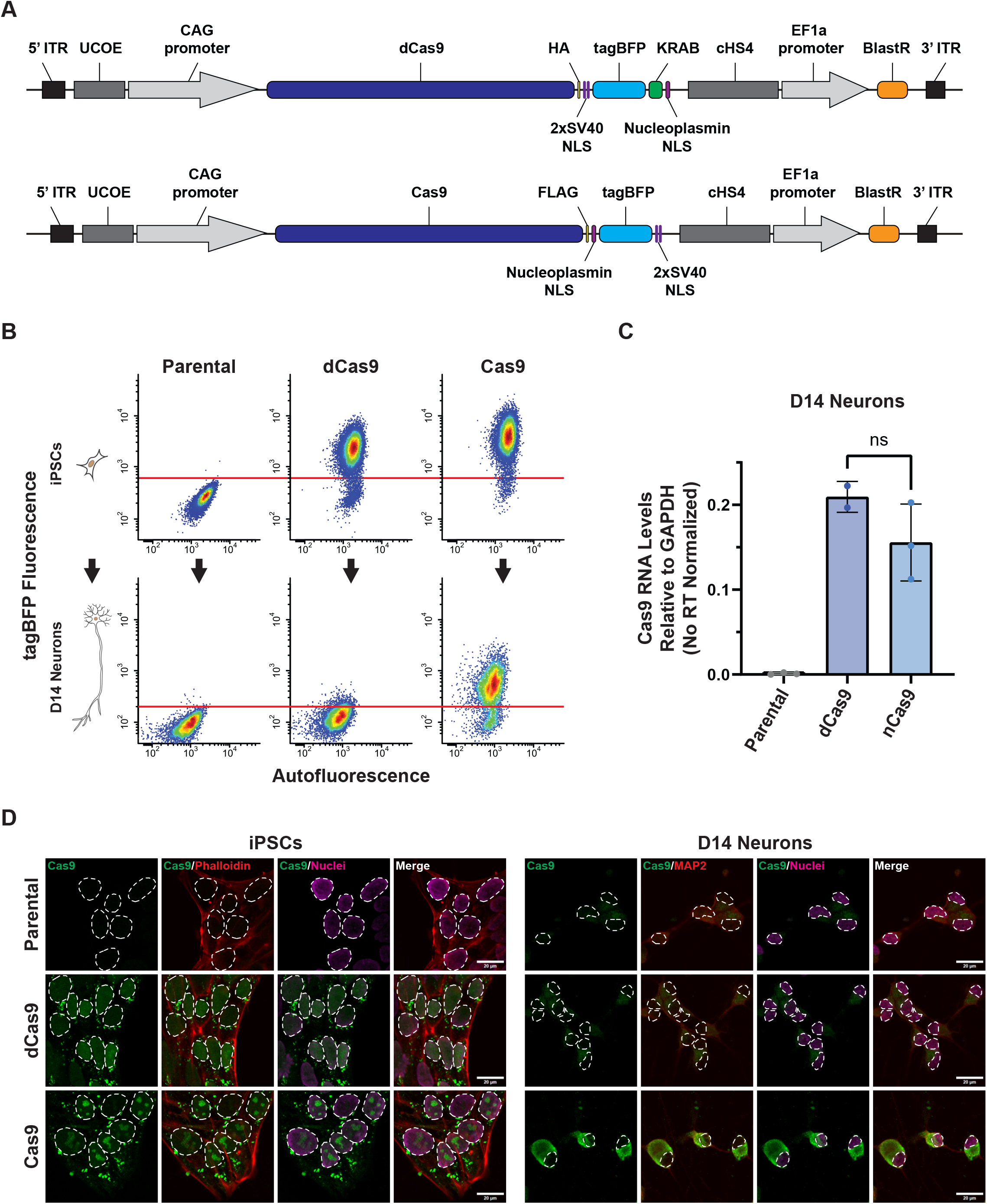
Mislocalization and differential protein stability of nCas9 and dCas9KRAB in iPSC-derived neurons post-differentiation. A - Illustrations of transposase donor plasmids for making stable dCas9 and nCas9 lines in KOLF2.1J_NGN2 iPSCs (drawn to scale). B - Flow cytometry plots measuring dCas9-KRAB or nCas9 expression via tagBFP fluorescence in stable iPSCs (top) vs expression in neurons 14 days post induction of differentiation with dox (bottom). Red line indicates the approximate cutoff for positive cells based on parental line autofluorescence. dCas9-KRAB and nCas9 expression are comparable at the iPSC stage; however, dCas9-KRAB neurons appear to lose almost all expression whereas the expression distribution of nCas9 neurons mirrors that of nCas9 iPSCs. Plots represent a minimum of 10,000 analyzed single cells for iPSCs and 3000 analyzed single cells for neurons. C - Quantification of dCas9-KRAB and nCas9 mRNA via RT-qPCR in Day 14 neurons shows no significant difference in gene expression across cell lines, suggesting that loss of dCas9-KRAB in neurons is independent of gene silencing. Values are shown relative to GAPDH expression after being normalized to no-RT samples. Data represents 2-3 independent wells per line. Error bars represent mean ± SD. Unpaired T-test with Welch’s correction was used to compare RNA levels. ns P=0.1694. D - Representative images of dCas9-KRAB and nCas9 protein localization in iPSCs (left) and Day 14 neurons (right) with ICC using a Cas9 antibody (green). Cells are stained for actin (phalloidin) or with a MAP2 antibody (red), and nuclei with NucSpot 750/780 (magenta). Nuclei are outlined for reference. In iPSCs, both versions show localization to the nucleus (dCas9-KRAB is more diffuse, whereas Cas9 appears more nucleolar), along with bright cytoplasmic accumulations. In D14 neurons, both versions show almost exclusively punctate cytoplasmic staining, similar to the cytoplasmic staining in iPSCs. Images are Max IPs of 3-plane (iPSCs) or 4-plane (neurons) Z-stacks (0.6um) taken at 40x with a spinning disk confocal. All images within cell type are adjusted to the same LUTS, based on background Cas9 antibody staining in the parental line. iPSCs and neurons are shown at the same scale; scale bars represent 20um.

To test if this was due to effect on the protein of mRNA, we performed RT-qPCR to measure dCas9-KRAB/nCas9 RNA levels in Day 14 neurons, using a primer set generic to both versions (Fig 1C, STAR methods). We observed no significant difference between the mRNA levels of the dCas9-KRAB and nCas9 in neurons suggesting that the observed effects on fluorescence is due to effects on the protein and not due to differential transgene silencing. This is also consistent with the fact that both cassettes have been introduced in the same donor backbone and displayed similar expression distributions in the iPSC state. To determine if there were any other differences in protein behavior between cell types, we performed immunofluorescence against dCas9-KRAB and nCas9 in both iPSCs and neurons using a single antibody against SpCas9 which recognizes both versions (Fig 1D). Staining in iPSCs showed both dCas9-KRAB and nCas9 in the nucleus. Interestingly, dCas9-KRAB appeared primarily diffuse while nCas9 had a predominantly nucleolar pattern, which could be due to differential interactions mediated by the KRAB domain. We also observed bright accumulations of both proteins in the cytoplasm, possibly related to effects of overexpressed proteins. In neurons, however, we observed that nCas9 was almost completely absent from the nucleus and restricted to the cytoplasm of the soma. Consistent with our flow data, dCas9 protein was almost undetectable by immunofluorescence (IF) as well, but appeared to also be restricted to bright puncta in the cytoplasm. Based on these observations, we hypothesized that neuronal-specific cytoplasmic mislocalization of the CRISPR machinery is leading to targeted degradation of dCas9-KRAB as the KRAB domain is an endogenous human protein domain derived from the *KOX1* transcription factor.

### Addition of a neuronal NLS improves nuclear localization and knockdown post-differentiation

To test the hypothesis that cytoplasmic mislocalization leads to KRAB-dependent degradation of CRISPR machinery in neurons, we generated a series of modified dCas9-KRAB piggybac constructs with an MeCP2 NLS and in combination with other alternative NLSs. We expected MeCP2 to be highly functional given it is derived from an essential neuronal nuclear protein and has previously been reported to be more effective in neurons(Karasev et al. 2022) (Fig 2A). Similarly to before, we generated stable polyclonal iPSC lines with these new constructs and compared them with our original dCas9-KRAB and nCas9 lines using flow cytometry in iPSCs (Fig S1A) and in day 14 neurons (Fig 2B). We found that the addition of an MeCP2 NLS was sufficient to rescue expression of dCas9-KRAB to near-nCas9 levels in day 14 neurons and boosted iPSC expression as well. Other NLSs paired with the MeCP2 NLS seemed to provide little or no additive effect; however, the 2xMeCP2 NLS appeared to provide a further boost to expression. Consistent with our flow data, western blots of total protein samples from Day 14 neurons showed no significant difference between nCas9 and 2xMeCP2 NLS-dCas9-KRAB protein levels (Fig 2D).

**Figure 2.**
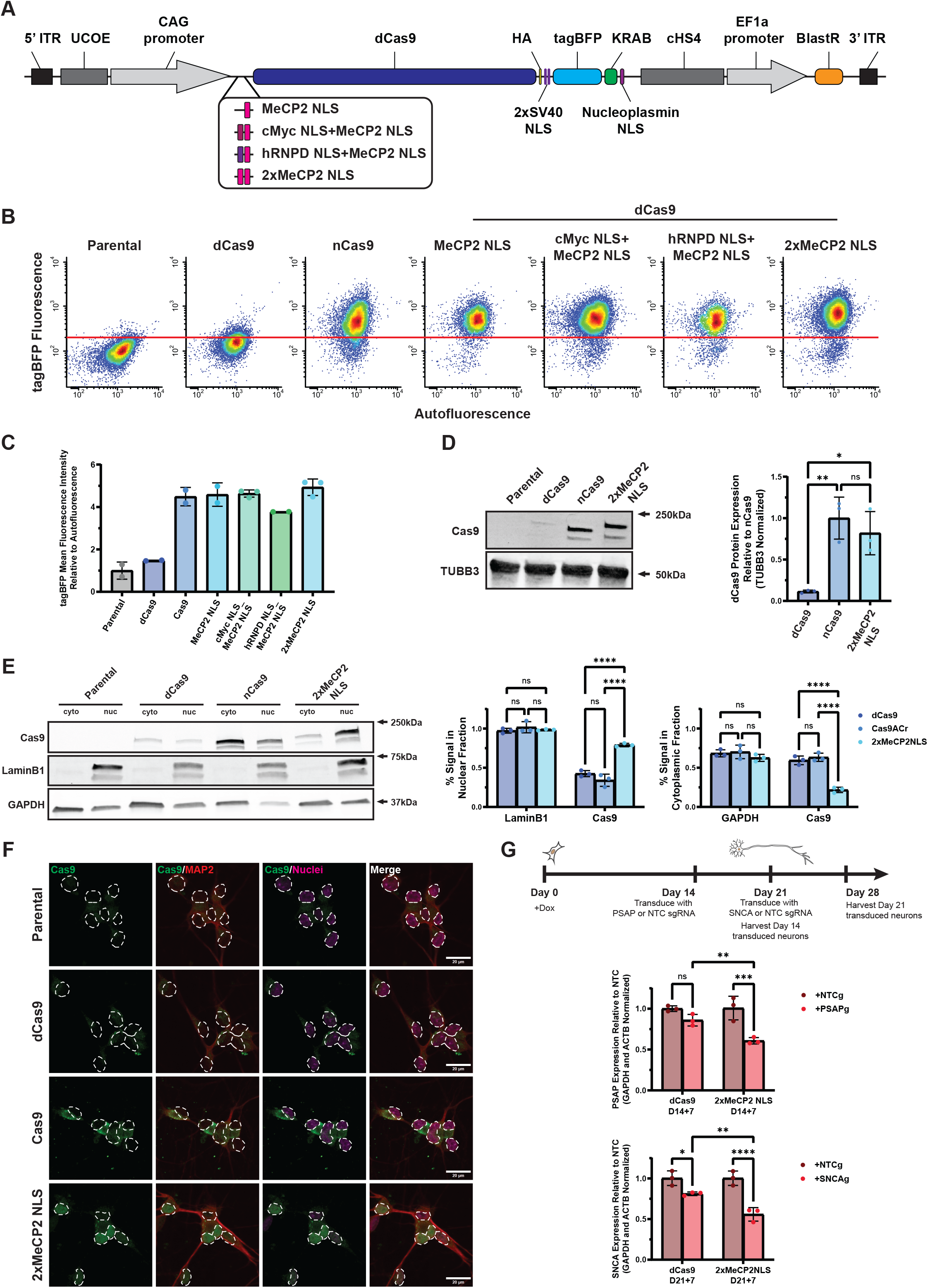
Alternative NLSs improve dCas9-KRAB nuclear localization in iPSC-derived neurons and stabilize protein levels during differentiation. A - Illustration of modified transposase donor plasmids to test the effect of alternative NLSs on dCas9-KRAB localization and stability in neurons (drawn to scale). B - Flow cytometry plots measuring expression of modified dCas9-KRAB constructs or nCas9 via tagBFP fluorescence in neurons 14 days post induction of differentiation with dox from stably integrated iPSCs. Red line indicates the approximate cutoff for positive cells based on parental line autofluorescence. Plots represent a minimum of 3000 analyzed single cells. C - Quantification of flow cytometry data showing the mean fluorescence intensity of tagBFP normalized to autofluorescence of dCas9-KRAB constructs and nCas9 in Day 14 neurons (Parental autofluorescence=1). Alternative NLS sequences rescue dCas9-KRAB expression to levels comparable to nCas9, with the 2xMeCP2 NLS version having the highest expression with a mean population intensity of 4.931x the parental, compared to a mean of 4.586x for the MeCP2 NLS construct and 4.636x for the cMyc NLS+MeCP2 NLS construct. Data represent the mean fluorescence intensity of 2-3 wells per line with a minimum of 2000 analyzed single cells per sample. Error bars represent mean ± SD. D - Representative Western blots (left) and quantification (right) of total protein from Day 14 neurons probed for Cas9 showing significant rescue of total protein levels of 2xMeCP2 NLS-dCas9-KRAB compared to the original dCas9-KRAB, with levels comparable to nCas9. Cas9 antibody signal was normalized to TUBB3 levels. Data represents 3 independent wells per line. Error bars represent mean ± SD. Protein levels between lines were compared with One-way ANOVA with Tukey’s multiple comparisons test. ^**^P=0.0050 for dCas9-KRAB vs Cas9, ^*^P=0.0150 for dCas9-KRAB vs 2xMeCP2 NLS-dCas9-KRAB, ns P=0.5719 for nCas9 vs 2xMeCP2 NLS-dCas9-KRAB. E-Representative Western blots (left) and quantifications (right) of dCas9-KRAB or nCas9 protein levels in nuclear and cytoplasmic protein fractions from Day 14 neurons. The original dCas9-KRAB and nCas9 are both predominantly cytoplasmic with nearly identical distributions despite much lower dCas9-KRAB levels. The 2xMeCP2 NLS-dCas9-KRAB construct shows a significant shift into the nuclear fraction which corresponds with the rescue of total protein levels. LaminB1 and GAPDH were used as nuclear and cytoplasmic controls, respectively, and were probed on replicate sample blots in parallel with Cas9. Quantifications for each protein represent the intensity of the signal in the respective fraction divided by the sum of intensities across both fractions. Cas9 quantifications were performed on respective LaminB1 and GAPDH blots. Data represent 3 independent wells per line. Error bars represent mean ± SD. Fractionation control and Cas9 antibody signals across cell lines were compared with 2-way ANOVA with Tukey’s multiple comparisons test. For Cas9 on LaminB1 blots: ns P=0.1236 for dCas9-KRAB vs nCas9, ^****^ P<0.0001 for dCas9-KRAB vs 2xMeCP2 NLS-dCas9-KRAB, ^****^ P<0.0001 for nCas9 vs 2xMeCP2 NLS-dCas9-KRAB. For Cas9 on GAPDH blots: ns P=0.6805 for dCas9-KRAB vs nCas9, ^****^ P<0.0001 for dCas9-KRAB vs 2xMeCP2 NLS-dCas9-KRAB, ^****^ P<0.0001 for nCas9 vs 2xMeCP2 NLS-dCas9-KRAB. F - Representative images of dCas9-KRAB protein localization in Day 14 neurons with ICC using a Cas9 antibody (green). Cells are stained with a MAP2 antibody (red) and nuclei with NucSpot 750/780 (magenta). Nuclei are outlined for reference. 2xMeCP2 NLS-dCas9-KRAB shows improved nuclear localization over original nCas9 and dCas9-KRAB constructs and higher intensity staining than the original dCas9-KRAB, consistent with the results from biochemistry assays. Images are Max IPs of 4-plane Z-stacks (0.6um) taken at 40x with a spinning disk confocal. All images are adjusted to the same LUTS, based on background Cas9 antibody staining in the Parental line. Scale bars represent 20um. G - Quantification of RT-qPCR data showing significantly improved knockdown of PSAP (top) and SNCA (bottom) in neurons expressing the 2xMeCP2 NLS-dCas9-KRAB vs the original dCas9-KRAB. Neurons were transduced with lentivirus expressing an mScarlet marker with either a targeting or non-targeting sgRNA at Day 14 (PSAP) or Day 21 (SNCA) post-differentiation and RNA harvested 7-days post-transduction. Gene expression levels were normalized to GAPDH and ACTB and are shown relative to the non-targeting average. Data represent 3 independent wells per transduction per line. 2-way ANOVA with uncorrected Fisher’s LSD was used to compare targeting guide with non-targeting guide within cell lines and targeting guides across cell lines. For PSAP KDs: ns P=0.0745 for dCas9-KRAB+NTCg vs +PSAPg, ^***^ P=0.0004 for 2xMeCP2 NLS-dCas9-KRAB+NTCg vs +PSAPg, ^**^ P=0.006 for dCas9-KRAB+PSAPg vs 2xMeCP2 NLS-dCas9-KRAB+PSAPg. For SNCA KDs: ^*^ P=0.0152 for dCas9-KRAB+NTCg vs +SNCAg, ^****^ P<0.0001 for 2xMeCP2 NLS-dCas9-KRAB+NTCg vs +SNCAg, ^**^ P=0.0035 for dCas9-KRAB+SNCAg vs 2xMeCP2 NLS-dCas9-KRAB+SNCAg.

We next tested whether the 2xMeCP2 NLS-dCas9-KRAB construct also displayed corrected nuclear localization using both biochemistry and imaging. We isolated cytoplasmic and nuclear protein fractions from Day 14 neurons using digitonin extraction of cytoplasmic protein followed by extraction of nuclear protein with Radioimmunoprecipitation Assay (RIPA) buffer (Fig 2E). We observed that our original dCas9-KRAB and nCas9 constructs, which share the same original NLSs, had nearly identical cytoplasmic/nuclear distributions, with the majority of the signal coming from the cytoplasmic fraction, confirming our previous imaging results (Fig 1D). The 2xMeCP2 NLS-dCas9-KRAB construct, however, showed a large and significant shift into the nuclear fraction, with a majority of the signal coming from the nuclear fraction. We then performed imaging to corroborate the fractionation results (Fig 2F). Similarly to before, the original dCas9-KRAB construct was nearly undetectable by IF, with only occasional cytoplasmic puncta visible, and the nCas9 was largely restricted to the cytoplasm as well. The 2xMeCP2 NLS-dCas9-KRAB construct, however, displayed an easily detectable and predominantly diffuse nuclear signal in most cells, at levels comparable to nCas9, suggesting that the corrected nuclear localization was able to rescue the dCas9-KRAB specific degradation. Lastly, we tested whether rescue of localization and expression would improve knockdown efficiency at the neuronal stage (Fig 2G). We lentivirally transduced sgRNAs in a fluorescent backbone at similar transduction levels) in either Day 14 (Fig. S1B) or Day 21 (Fig. S1C) neurons and harvested RNA 7 days post-transduction. In both cases, we observed increased knockdown in the 2xMeCP2 NLS neurons compared to the original dCas9-KRAB line, suggesting that the corrected construct can improve sgRNA knockdown efficiency.

To test whether the defects we observed in nuclear localization and protein expression in neurons were construct-specific or common to any variations of CRISPR machinery, we tested an alternative CRISPRi design, iE61 PB-Zim3-XTEN-dCas9-mScarlet-puro-BFP, which uses the *ZIM3* domain instead of the *KOX1* KRAB domain, but similarly relies on SV40 NLSs for nuclear localization(Alerasool et al. 2020). Imaging confirmed that this alternative CRISPRi design similarly failed to effectively localize to the nucleus of neurons, suggesting that SV40 NLSs are generally ineffective in neurons for CRISPR constructs (Fig. S2A). Interestingly, despite the lack of nuclear localization, the Zim3-dCas9 appeared to retain stable expression in neurons unlike mislocalized dCas9-KRAB (Fig. S2B, S2C). This supports the hypothesis that the loss of expression of cytoplasmic-restricted dCas9-KRAB is mediated by the KRAB domain and is specific to the commonly used *KOX1* domain.

## Discussion

The use of CRISPR enzymes in postmitotic, iPSC-derived cell lines holds immense promise for disease modeling, therapeutic development and modifier screenings. High throughput CRISPR-based screening methods in iPSC-derived neurons and glia cells have already been applied successfully for elucidating both disease and basic biology(Tian et al. 2021; Tian et al. 2019; Parra Bravo et al. 2024), but have also been limited by the constraints of ensuring activity of sgRNAs and CRISPR machinery post-differentiation. To limit the inefficiencies associated with the delivery of either or both components at the neuron-stage, these studies have relied on delivery of these components in iPSCs followed by differentiation.

Here, using piggybac delivery of CRISPR machinery to avoid the need for iPSC clonal engineering, we found a surprising discrepancy in the stability of nCas9 and dCas9-KRAB expression in iPSC-derived neurons which was not explained by differences in mRNA levels. This led us to find that both dCas9 and nCas9 failed to efficiently localize to the nucleus in neurons despite having a robust nuclear presence in iPSCs. We hypothesized that this mislocalization may be responsible for the dCas9-KRAB-specific repression we observed. To test this, we determined whether alternative, neuronal specific NLSs, would rescue these dCas9-KRAB deficits in neurons. We indeed found that the addition of two tandem MeCP2 NLSs was able to rescue both neuronal dCas9-KRAB protein levels and drastically improve nuclear localization. We additionally showed that this rescue was sufficient to improve the knockdown efficacy of guides delivered directly to mature neuronal cultures.

While the exact mechanism of KRAB domain-mediated protein loss in neurons remains unclear, it seems likely that its mislocalization to the cytoplasm engages targeted protein degradation mechanisms. Given that the KRAB domain is derived from a human transcription factor, specifically here *KOX1*, the most commonly used version, it may be subject to endogenous regulatory pathways that would recognize and degrade the protein when not in the nucleus or not bound to DNA. On the other hand, nCas9 is a totally ectopic protein, so its “mislocalization” to the cytoplasm is unlikely to be recognized by cells. It also appears that this degradation is not a neuron-specific pathway, as we saw that our improved NLS constructs boosted dCas9-KRAB expression in iPSCs as well. Rather, the neuron-specific deficit in nuclear localization amplified the effect to the point of near-total protein loss.

Importantly, while our studies were performed in iPSC-derived neurons, we expect that other differentiated, post-mitotic cell types may suffer from similar deficits in both localization of CRISPR constructs and mislocalization-dependent loss of dCas9-KRAB expression. Furthermore, we expect all SV40 NLS-dependent CRISPR enzymes, including those for CRISPRa and prime editing, to suffer from these localization deficits, whereas resulting changes in protein stability will be construct-dependent. It is possible that even in dividing cells, the SV40 NLS does not deliver Cas9 into the nucleus through the nuclear pore, but rather it localizes to the nucleus during cell division due to Cas9 or KRAB affinity to DNA. The limited nuclear expression of CRISPR enzymes in postmitotic cells may have dramatic implications on the efficiency of gene perturbation and editing beyond iPSC-derived neurons in both cultured cells and in-vivo for research and therapeutic development and will need to be further investigated.

Lastly, implementation of effective, post-differentiation gene perturbation technologies will require overcoming additional challenges including optimization of transgene expression, cell type-specific epigenetic landscapes and a wide range of protein and mRNA stabilities in post-mitotic cells. Our work here represents one important step forward in maximizing the effectiveness of CRISPR-based technologies in iPSC-derived neurons, and has significant implications for other differentiated cell types and therapeutic development. While additional work may be required to ensure machinery can be delivered and/or expressed at saturating levels and to ensure that employed sgRNAs are optimal for targeting the neuronal genome, robust nuclear levels of CRISPR machinery are a necessary prerequisite for all further steps in applying these techniques.

*Figure S1* - NLS-mediated improvements in dCas9-KRAB expression are not neuron-specific and allow for improved knockdown efficiency after differentiation.

A - Flow cytometry histograms of stable iPSCs expressing the original dCas9-KRAB and nCas9 constructs and the dCas9-KRAB constructs with alternative NLSs. The MeCP2 NLS increased dCas9-KRAB expression and the 2xMeCP2 NLS construct further boosted levels to match nCas9 levels. Other NLSs in combination with a single MeCP2 NLS provided no additive benefit to dCas9-KRAB expression.

B and C-Flow cytometry histograms of neurons 7 days after lentiviral transduction with either a non-targeting control guide and a guide targeting PSAP at Day 14 (BA) or a non-targeting control guide and a guide targeting SNCA at Day 21 (C). Based on mScarlet fluorescence, almost all neurons were transduced. Expression distributions for each guide at respective timepoints were comparable between dCas9-KRAB and 2xMeCP2 NLS-dCas9-KRAB neurons, so differential knockdown efficiencies between lines are likely not due to differences in transduction and sgRNA expression.

Figure S2 - Nuclear localization defects are common to SV40NLS-dependent CRISPR constructs while mislocalization-dependent expression loss is KRAB domain-specific.

A - Representative images of Zim3-dCas9 protein localization in Day 14 neurons with ICC using a Cas9 antibody (green). Cells are stained with a MAP2 antibody (red) and nuclei with NucSpot 750/780 (magenta). Nuclei are outlined for reference. Despite showing stable protein expression, this alternative construct, which also relies onSV40 NLSs, shows a similar defect in nuclear localization in neurons. Images are a single widefield plane taken at 40x. All images are adjusted to the same LUTS. Scale bars represent 20um.

B - Representative Western blots of total protein from Day 14 neurons probed for Cas9 showing rescue of total protein levels with the addition of an MeCP2 NLS. We also show that an alternative dCas9 using the Zim3 domain instead of the KOX1 KRAB domain also retains stable protein expression during differentiation. C - Quantification of dCas9 levels from western blots. Cas9 antibody signal was normalized to TUBB3 levels. Data represents 3 independent wells per line. Error bars represent mean ± SD.

## Acknowledgements

We would like to thank all members of the Shalem lab for support and discussions related to this project and Michale Ward from NIH/NINDS for providing piggybac backbone vectors and for discussions related to this work. Funding for this project came from NIH/NIGMS grant DP2-GM137416, TargetALS grant and CZI pilot grant all awarded to O.S.

## Methods

### Cloning

Original piggybac donor vectors were obtained from VectorBuilder. Guide cloning was performed via Golden Gate assembly in a CROPseq_v2 backbone modified to express 2xcMycNLS-mScarlet-P2A/T2A-puromycin. All oligos for cloning were purchased from IDT. PCRs were performed with PrimeSTAR® GXL DNA Polymerase (Takara, R050A). Custom plasmids were generated with NEBuilder® HiFi DNA Assembly Master Mix (NEB, E2621S).

iE61 PB-Zim3-XTEN-dCas9-mScarlet-puro-BFP was a gift from iPSC Neurodegenerative Disease Initiative (iNDI) & Michael Ward (Addgene plasmid # 204722; http://n2t.net/addgene:204722; RRID:Addgene_204722)

### iPSC Maintenance

KOLF2.1J_NGN2 iPSCs (The Jackson Laboratory), described previously (Pantazis et al., 2022; Reilly et al., 2023), were cultured in mTesR Plus medium (STEMCELL Tech, 100-0276) on 6-well cell culture plates coated with hESC-Qualified, LDEV-Free, Matrigel Matrix (Corning, 354277) diluted according to manufacturer’s lot number recommendation. Media was replaced every other day until 80%–90% confluent when cells were then passaged with Versene (ThermoFisher Scientific, 15040066). Media was aspirated and cells washed 2x with DPBS (ThermoFisher Scientific, 14190144) and then incubated with Versene at 37C for 5-7min. Versene was then aspirated, and cells lifted by washing the well with fresh mTesR Plus medium and gently scraping if needed. Colonies were broken up by gently triturating the cell mixture before transferring cells to a new Matrigel-coated plate at desired concentration.

### Generation of stable iPSC lines

iPSCs were collected using Accutase (ThermoFisher Scientific, A1110501) by aspirating media, washing 2x with DPBS, and then incubating with Accutase at 37C for 10min. Accutase was then diluted with mTesR Plus medium supplemented with 10nM Y-27632 dihydrochloride ROCK inhibitor (Tocris, 125410) and iPSCs pelleted and then resuspended in fresh mTeSR Plus with Y-27632 for counting. iPSCs were plated at 1.5×10^6^ cells per well of a 6-well plate on hESC Matrigel in 1.5mL of media. After at least 1 hour, iPSCs were transfected using Lipofectamine™ Stem Transfection Reagent (ThermoFisher Scientific, STEM00008) at 5uL per well with 1ug of DNA at a donor to transposase ratio of 2:1. The following day, iPSCs were split again with Accutase and then maintained in mTeSR Plus with Y-27632 and Blasticidin S HCl (ThermoFisher Scientific, A1113903) selection at 10ug/mL for at least 1 week prior to cell sorting, with Accutase passaging as needed. To generate polyclonal stable lines, the top 50% of cells with distinct tagBFP signal were kept with a minimum of 10,000 cells sorted for any given line. Following sorting, iPSCs were allowed to recover in mTeSR Plus with Y-27632, blasticidin selection, and Penicillin-Streptomycin (ThermoFisher Scientific, 15070063). Y-27632 was removed 48-72 hours post-sorting, and iPSCs were maintained as normal with Versene passaging as needed. Blasticidin and Pen-Strep were removed 1 week post-sorting.

### Neuronal Differentiation

KOLF2.1J_NGN2 iPSCs were differentiated using doxycycline-induced expression of NGN2 based on previously described methods (Pantazis et al., 2022; Reilly et al., 2023). Day 0 iPSCs were collected using Accutase (ThermoFisher Scientific, A1110501) by aspirating media, washing 2x with DPBS, and then incubating with Accutase at 37C for 10min. Accutase was then diluted with mTesR Plus medium supplemented with 10nM Y-27632 dihydrochloride ROCK inhibitor (Tocris, 125410) and iPSCs pelleted and then resuspended in fresh mTeSR Plus with Y-27632 for counting. iPSCs were plated in Pre-Differentiation Medium, comprised of KnockOut™ DMEM/F-12 (ThermoFisher Scientific, 12660012), 1x N2 supplement (ThermoFisher Scientific, A1370701), 1X MEM Non-Essential Amino Acids (ThermoFisher Scientific, 11140-050), and 1X GlutaMAX Supplement (ThermoFisher Scientific, 35050-061), supplemented with 10nM Y-27632 and 2ug/mL doxycycline hydrochloride (Sigma-Aldrich, D3072). iPSCs were plated at a concentration of 1×10^6^ cells/well on 6-well plates coated with Matrigel, Growth Factor Reduced Basement Membrane Matrix, LDEV-free (Corning, 354230), diluted to 0.5 mg/plate. Media was changed daily. After Day 0, Y-27632 was removed. On Day 3, 5-fluoro-20-deoxyuridine and uridine (Sigma-Aldrich, F0503 and U3003) were added at 10uM each. On Day 4, pre-differentiated cells were collected with Accutase as above and resuspended for counting and replating in Maturation Media, comprised of Neurobasal Plus Medium (ThermoFisher Scientific, A3582901), 1x B27 Plus supplement (ThermoFisher Scientific, A3582801), 1x CultureONE supplement (ThermoFisher Scientific, A3320201), 1X MEM Non-Essential Amino Acids, and 1X GlutaMAX, supplemented with with 10nM Y-27632, 2ug/mL doxycycline hydrochloride, 10ng/mL BDNF (R&D, 248-BDB/CF), 10ng/mL NT-3 (PeproTech, 450-03), 10ng/mL GDNF (R&D, 212-GD\CF) and 200μM L-ascorbic acid (Sigma-Aldrich, A8960). 12-well plates or 24-well glass-bottomed dishes (Cellvis, P24-1.5H-N) were prepared for replating by coating with 100 ug/mL poly-L-ornithine (Sigma-Aldrich, P3655) overnight at 37C, washing 3x with H2O, and drying overnight at room temperature. Prior to coating, glass-bottomed plates were pre-treated with 1.0N HCl (Sigma-Aldrich, H9892) for at least 15min and washed 1x with DPBS and then 2x with H2O. Prior to replating, coated and dried plates were pre-incubated with plain Neurobasal Plus media at 37C while cells were prepared for replating. Pre-incubation media was aspirated, and pre-differentiated cells plated at 1×10^5^ cells/well for 12-well plates for flow cytometry and at 5×10^4^ cells/well on 24-well plates for imaging. After 1hr, at which point cells are well-adhered, an additional volume of supplemented Maturation Media was added, but without Y-27632 and with laminin (ThermoFisher Scientific, 23017015) at 2ug/mL (final concentration per well of 1ug/mL). Thereafter, half media changes were performed 1-2x per week with supplemented Maturation Medium without Y-27632 or doxycycline and with laminin at 1ug/mL.

### Flow cytometry analysis of neurons and iPSCs

Neurons were dissociated for flow cytometry analysis using papain (Worthington, LK003176) and resuspended in 0.5mL of base Maturation Medium with 10nM Y-27632, while iPSCs were dissociated with Accutase and resuspended in mTeSR Plus with 10nM Y-27632. Cells were kept on ice and passed through a 40µm cell strainer prior to flow analysis and/or sorting for iPSCs. Cellular fluorescence was measured on a BD FACSAria Fusion (BD Biosciences) using an 85µm nozzle. For dCas9/nCas9 measurements, mtagBFP fluorescence was detected by the 405nm laser and the 450/50 filter and autofluorescence was detected by the 561nm laser and the 582/15 filter. For viral titering, mScarlet fluorescence was measured by the 561nm laser and filters 600LP and 610/20. Data were analyzed using the R package CytoExploreR (v1.1.0).

### IF and confocal microscopy of neurons and iPSCs

Neurons and iPSCs grown in glass-bottomed dishes were washed 3x with PHEM Buffer (Electron Microscopy Sciences, 11163), and fixed in 4% formaldehyde plus 0.25% glutaraldehyde (Electron Microscopy Sciences, 15710 and 16120) in PHEM buffer. Fixed cells were washed 3x in DPBS and then blocked and permeabilized in 5% goat serum (Cell Signaling Technology, 5425) and either 0.25% TritonX-100 (VWR, 0694) in DPBS for neurons or 0.1% TritonX-100 for iPSCs. Cells were incubated with primary antibodies diluted in blocking buffer overnight at 4C. Cells were then washed 3x with DPBS for 5min each and then incubated with secondary antibodies or cell stains diluted in blocking buffer for 1hr at room temperature. Cells were washed 1x with DPBS for 5min, then incubated with NucSpot 750/780 (Biotium, 41038) diluted at 1:10,000 in DPBS for 5min, followed by 1x DPBS wash for 5min. Cells were finally stored in ibidi Mounting Medium (ibidi, 50001) at 4C. Images were acquired on a Leica TCS SP8 confocal microscope. Z-stacks (0.6μm slices) were recorded with 40× Plan-Apochromat lenses, 1.4 NA. The following primary antibodies were used: Cas9 (Takara, 632607; 1:150), MAP2 (Abcam, ab5392; 1:5000). The following secondary antibodies and cell stains were used: Goat anti-Rabbit IgG (H+L) Highly Cross-Adsorbed Secondary Antibody, Alexa Fluor™ Plus 488 (ThermoFisher Scientific, A32731; 1:2000), Goat anti-Rabbit IgG (H+L) Highly Cross-Adsorbed Secondary Antibody, Alexa Fluor™ 555 (ThermoFisher Scientific, A21429; 1:2000), Goat anti-Chicken IgY (H+L) Cross-Adsorbed Secondary Antibody, Alexa Fluor™ Plus 647 (ThermoFisher Scientific, A32933; 1:1000), Alexa Fluor™ 647 Phalloidin (ThermoFisher Scientific, A22287; 1:1000).

### Total protein harvesting and neuronal fractionation

For total protein, neurons were washed 3x with cold DPBS+/+ (ThermoFisher Scientific, 14040133) and lysed in-well for 10min at 4C while rocking with 1x RIPA lysis buffer (Cell Signaling, 9806) plus 1x protease inhibitor cocktail (Sigma-Aldrich, P8340).

To extract cytoplasmic protein, neurons were washed 3x with cold DPBS+/+ and then treated in-well for 10min at 4C while rocking with 0.05% Digitonin (ThermoFisher Scientific, BN2006) diluted in DPBS with 1x protease inhibitor cocktail (PIC). Cytoplasmic lysate was then collected and RIPA+PIC added to final 1x concentrations. Neurons were then washed 1x with cold DPBS+/+ and finally lysed with 1x RIPA + 1x PIC as for total protein to extract remaining nuclei.

After lysis, all samples were sonicated for 30s and then centrifuged for 10min at 14000g at 4C. Supernatants were transferred to fresh tubes and stored at -80C prior to denaturation.

### Western Blotting

Protein samples were denatured at 95C for 5min in 1x Laemmli buffer (Bio-Rad, 1610747) with 2.5% 2-mercaptoethanol (Sigma-Aldrich, M6250) and then loaded on precast stain-free SDS-PAGE gels (Bio-Rad, 4568094 and 4568096). Immunoblotting followed using standard protocols with Immun-Blot® Low Fluorescence PVDF membranes (BioRad, 1620260). 5% milk in 1x TBS with 0.1% Tween 20 (VWR, 0777) was used for blocking and antibody dilution. SDS (ThermoFisher Scientific, 15553027) was added to secondary antibody dilutions at a final concentration of 0.02%. Imaging of blots was performed on a LI-COR Odyssey instrument. Quantification of blots was performed using densitometry in ImageJ. The following primary antibodies were used: Cas9 (Cell Signaling, 14697; 1:1000), TUBB3 (Abcam, ab18207; 1:800), LaminB1 (Proteintech, 12987-1-AP; 1:2000), GAPDH (Cell Signaling, 2118; 1:500). The following secondary antibodies were used: IRDye 680LT Goat anti-Rabbit (LI-COR, 926-68021; 1:10,000), IRDye 800CW Goat anti-Mouse (LI-COR, 926-32210; 1:10,000).

### Transfection for lentiviral production

Lentivirus was produced from HEK293T cells seeded into gelatin-coated 6-well tissue culture dishes at 5×10^5^ cells/well. 24 hours after plating, transfection was performed with PEI (Polysciences, 24765) in Opti-MEM (ThermoFisher Scientific, 31985062). 100uL of Opti-MEM was incubated with 1) 1.06ug pMDLg 2) 0.57ug pMD2.G 3) 0.4ug pRSV-Rev 4) 1.06ug donor plasmid. 7.35 μl PEI was added dropwise to the DNA+Opti-MEM solution. The transfection mix was incubated for 15 minutes and then added dropwise to cells containing 2mL of antibiotic-free media. Media was removed 4-6 hours post-transfection, cells washed 1x with DPBS, and 2mL of base Maturation Medium added. Virus was collected 48 hours post-transfection and filtered through 0.45 μm cellulose acetate filters (VWR, 76479-040) or 0.45 μm PES filters (ThermoFisher Scientific, 50-607-518). Virus used within three days was stored at 4C; for longer-term storage, aliquots were frozen at -80C.

### Lentiviral transduction of neurons

Neurons were transduced at either Day 14 or Day 21 with lentivirus containing sgRNAs against target genes. Transduction was performed at the time of regular half-media changes. Virus prepared in Maturation Medium was added at 10% of final well volume (200uL per well for a 12-well) to fresh Maturation Medium followed by standard media supplementation. Approximately half of the media volume of each well was removed (accounting for evaporation) and then replenished with the respective media and virus solution. Following transductions, cells were maintained for 7 days before harvesting RNA.

### RNA isolation and qPCR

Total RNA was isolated from neurons using the Qiagen RNeasy Plus Mini kit (Qiagen, 74134) according to manufacturer’s recommendations, including the addition of 2-mercaptoethanol. cDNA was prepared using either the SuperScript™ IV VILO™ Master Mix with ezDnase (ThermoFisher Scientific, 11766050) for Cas9 gene expression experiments or the High-Capacity cDNA Reverse Transcription Kit (Applied Biosystems, 4368814) for knockdown experiments. qPCR analysis was performed with Power SYBR Green master mix (Applied Biosystems, 4367659) on a CFX384 Touch Real-Time PCR Detection System (Bio-Rad).

For evaluating dCas9/nCas9 expression in neurons, background signal from Cas9 primers in no RT controls for each sample was subtracted from respective RT samples, before normalizing RT sample signal to GAPDH.

For knockdown experiments, RNA levels for genes of interest were determined using the ΔΔCt method, using the geometric mean of ACTB and GAPDH as control genes.

Primers:

GAPDH F GGAGCGAGATCCCTCCAAAAT

GAPDH R GGCTGTTGTCATACTTCTCATGG

ACTB F ACCTTCTACAATGAGCTGCG

ACTB R CCTGGATAGCAACGTACATGG

Cas9 F GGAAGTTCGACAATCTGACCAAGG

Cas9 R TGCCACGTGCTTTGTGATCTG

PSAP F CCCGGTCCTTGGACTGAAAG

PSAP R TATGTCGCAGGGAAGGGATTT

SNCA F AAGAGGGTGTTCTCTATGTAGGC

SNCA R GCTCCTCCAACATTTGTCACTT

