## Supplementary figures and images for "CRISPR associated enzymes are mislocalized to the cytoplasm in iPSC-derived neurons resulting in KRAB-specific degradation"

### Figure S1

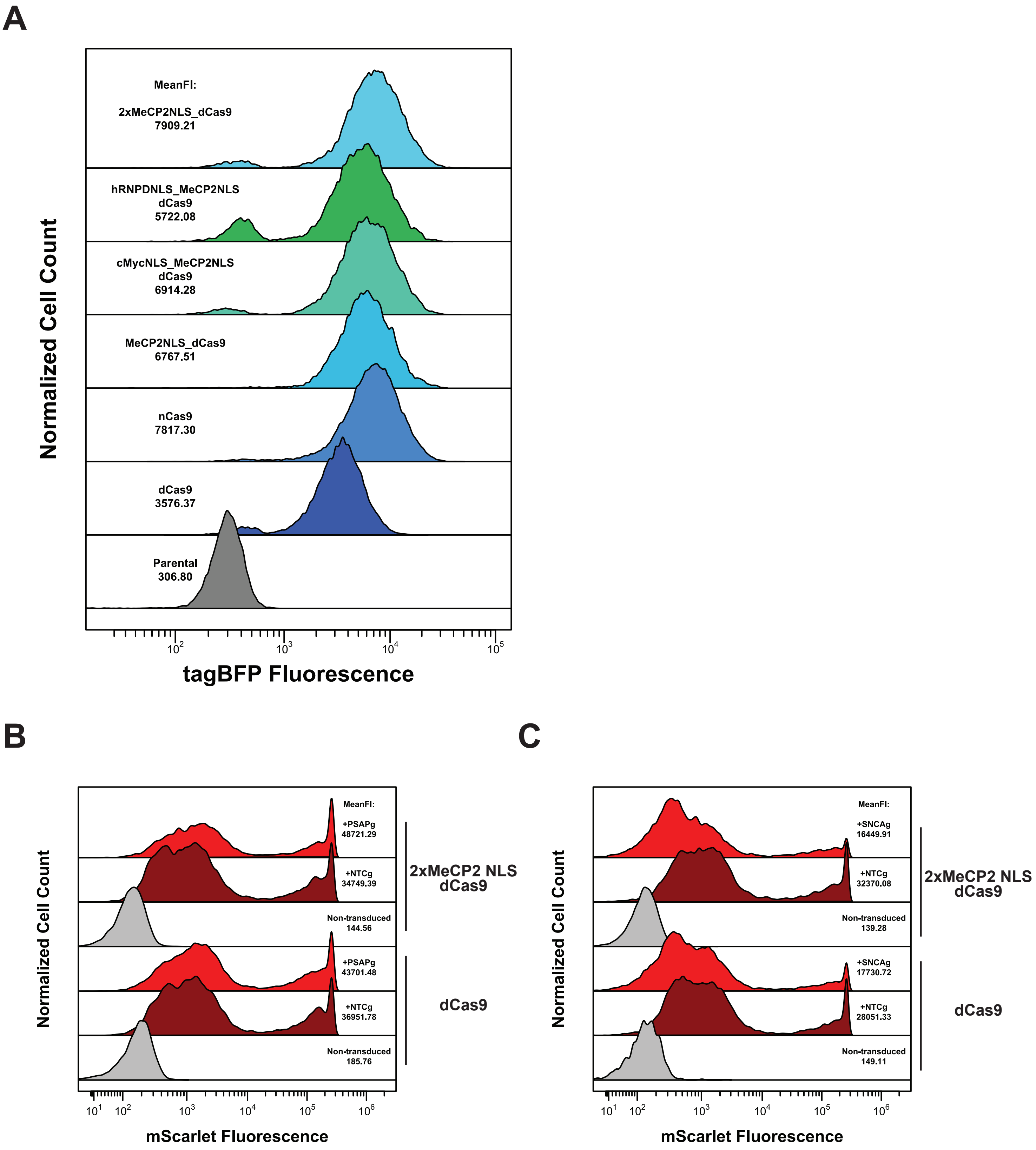

### Figure S2

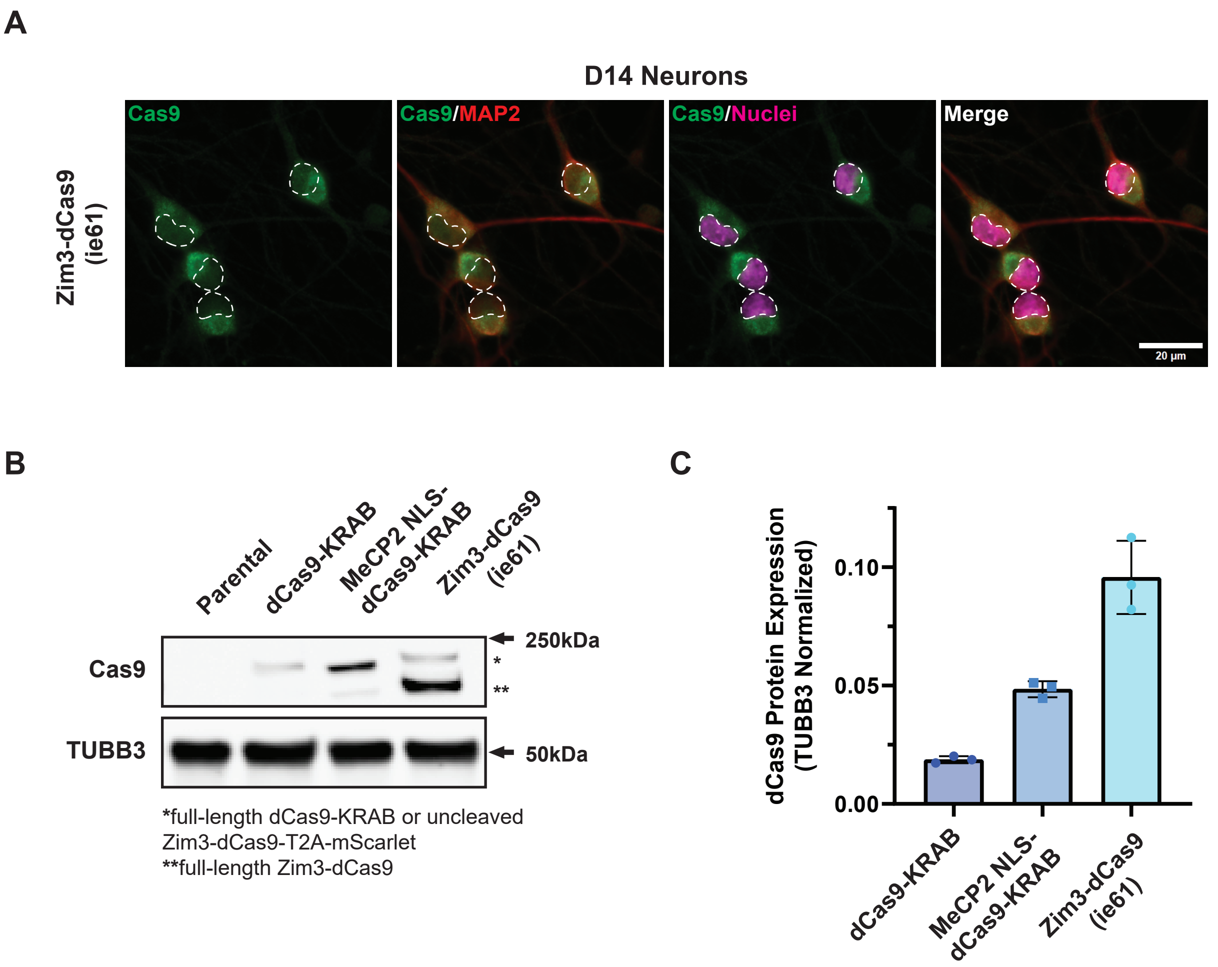
